# A Novel, Robust, and Portable Platform for Magnetoencephalography using Optically Pumped Magnetometers

**DOI:** 10.1101/2024.03.06.583313

**Authors:** Holly Schofield, Ryan M. Hill, Odile Feys, Niall Holmes, James Osborne, Cody Doyle, David Bobela, Pierre Corvilian, Vincent Wens, Lukas Rier, Richard Bowtell, Maxime Ferez, Karen J. Mullinger, Sebastian Coleman, Natalie Rhodes, Molly Rea, Zoe Tanner, Elena Boto, Xavier de Tiège, Vishal Shah, Matthew J. Brookes

## Abstract

Magnetoencephalography (MEG) measures brain function via assessment of magnetic fields generated by neural currents. Conventional MEG uses superconducting sensors, which place significant limitations on performance, practicality, and deployment; however, the field has been revolutionised in recent years by the introduction of optically-pumped-magnetometers (OPMs). OPMs enable measurement of the MEG signal without cryogenics, and consequently the conception of ‘OPM-MEG’ systems which ostensibly allow increased sensitivity and resolution, lifespan compliance, free subject movement, and lower cost. However, OPM-MEG remains in its infancy with limitations on both sensor and system design. Here, we report a new OPM-MEG design with miniaturised and integrated electronic control, a high level of portability, and improved sensor dynamic range (arguably the biggest limitation of existing instrumentation). We show that this system produces equivalent measures when compared to an established instrument; specifically, when measuring task-induced beta-band, gamma-band and evoked neuro-electrical responses, source localisations from the two systems were highly comparable and temporal correlation was >0.7 at the individual level and >0.9 for groups. Using an electromagnetic phantom, we demonstrate improved dynamic range by running the system in background fields up to 8 nT. We show that the system is effective in gathering data during free movement (including a sitting-to-standing paradigm) and that it is compatible with simultaneous electroencephalography (EEG – the clinical standard). Finally, we demonstrate portability by moving the system between two laboratories. Overall, our new system is shown to be a significant step forward for OPM-MEG technology and offers an attractive platform for next generation functional medical imaging.

## INTRODUCTION

Magnetoencephalography (MEG) measures the magnetic fields generated above the scalp by current flow through neuronal assemblies in the brain (Cohen, 1968). Mathematical modelling of these fields results in three dimensional images showing the spatial and temporal signatures of electrophysiological activity. MEG is a proven tool to investigate brain function, with applications in neuroscience and clinical practice (Baillet, 2017). In neuroscience, it can be used to measure evoked responses, neural oscillations, functional connectivity, and network dynamics – showing how the brain continually forms and dissolves networks in support of cognition. Clinically, MEG is most often used in epilepsy to localise the brain areas responsible for seizures as well as surrounding eloquent cortex (De Tiège et al., 2017). There are other potential applications ranging from study of diseases that are common in childhood (e.g., measurement of the auditory evoked response latency in autism (Matsuzaki et al., 2019)) to investigation of neurodegenerative conditions in older adults (e.g., the measurement of cortical slowing in dementia (Gouw et al., 2021)). MEG outperforms the clinical standard – electroencephalography (EEG) – in terms of both spatial precision (since magnetic fields are less distorted by the skull than the electric potentials measured by EEG) and sensitivity (since EEG is more effected by artefacts from non-neuronal sources – such as muscles) (Boto et al., 2019; Goldenholz et al., 2009). However, conventional MEG systems are based on cryogenically cooled (superconducting) sensors; this means systems have a high cost and are impractical for many applications, particularly compared to EEG. This has prevented widespread uptake of MEG systems.

In recent years, MEG instrumentation has been revolutionised by the introduction of optically pumped magnetometers (OPMs). (See (Brookes et al., 2022; Schofield et al., 2023; Tierney et al., 2019) for reviews.) OPMs measure magnetic fields with a similar sensitivity to the sensors used for conventional MEG, but without the need for cryogenic cooling. They can also be microfabricated (Schwindt et al., 2007; V. Shah et al., 2007, 2020; V. K. Shah & Wakai, 2013) such that they are small and lightweight. This leads to multiple advantages. For example, sensors can be sited closer to the scalp surface (compared to cryogenic devices, as a thermally insulating gap is no longer required); this improves signal amplitude significantly (Boto et al., 2016, 2017; Iivanainen et al., 2017, 2019, 2020) and theoretical calculations suggest this can offer unprecedented spatial resolution (higher than conventional MEG and EEG) (Nugent et al., 2022; Tierney et al., 2022; Wens, 2023). Arrays can be adapted to fit any head shape – from newborns to adults (Corvilain et al., 2024; Feys, et al., 2023; Hill et al., 2019; Rier et al., 2024). Adaptability also means arrays can be designed to optimise sensitivity to specific effects (Hill et al., 2024) or brain areas (Lin et al., 2019; Tierney, Levy, et al., 2021). As the sensors move with the head, participants can move freely during recordings (assuming background fields are well controlled) (Holmes et al., 2018, 2019, 2023; Rea et al., 2021). This enables the recording of data during novel tasks (Boto et al., 2018; Rea et al., 2022) or even epileptic seizures (Feys, et al., 2023; Hillebrand et al., 2023). The adaptability to different head size/shape coupled with motion robustness (Feys & De Tiège, 2024) mean that, like EEG, OPM-MEG systems arewearable. However, unlike EEG, sensors do not need to make electrical contact with the head, making OPM-MEG considerably more practical than EEG in terms of both patient friendliness and set up time. Finally, even at this early stage of development, OPM-based systems are becoming cheaper to buy and run than conventional MEG devices. These significant advantages could – in theory – lead to OPM-MEG becoming the method of choice for electrophysiological measurement, potentially even replacing EEG as a clinical tool for some applications.

Despite its promise, OPM-MEG remains in its infancy. The optimum system design has not yet been settled and the OPM sensors themselves remain limited in performance with higher noise compared to cryogenic sensors, lower bandwidth (though it is adequate for most MEG signals of interest) and much smaller dynamic range. To date, most published OPM-MEG studies have used systems composed of multiple independent sensors which are joined together and synchronised to form an array. Such systems work (e.g., (Boto et al., 2018; Corvilain et al., 2024; Feys et al., 2022; Hill et al., 2020; Iivanainen et al., 2019)) but their electronic architecture is complex and can be prone to failure. In addition, whilst the OPM-MEG helmet is lightweight, electronics racks are large, cumbersome, and must be kept outside the magnetically shielded room to limit magnetic interference. As a result, long cables must pass through waveguides in the MSR to the electronics. Such cabling can be prone to interference. Moreover, when systems have large sensor counts, cabling becomes cumbersome with large numbers of wires trailing from the subject. While this is fine for static systems, for systems where the aim is to allow the subject to move freely (even to walk around a room (Holmes et al., 2023)), having heavy cabling draped around a participant is impractical – particularly in the case of patients.

At a technical level, OPMs already have high sensitivity; indeed, even though noise levels are higher than conventional sensors (Boto et al., 2022), closer proximity to the scalp enables improved signal strength (Hill et al., 2024). However, perhaps their biggest limitation is dynamic range. This is because the sensor output is only a linear function of local magnetic field within a very narrow range – approximately −1.5 nT to +1.5 nT for rubidium OPMs. (For this reason, OPMs designed for MEG are often referred to as “zero-field” sensors, as they must operate in a ‘close to zero’ field.) Such a narrow range is problematic; whilst the MEG signal itself is small relative to this window, even in magnetically shielded environments the environmental field fluctuations can be much larger. The problem is even more complicated if the head is allowed to move, since even in the absence of a time-varying environmental fields, movement in a static field can take sensors outside their dynamic range. This problem is notionally solved by operating sensors in a “closed-loop”, mode, whereby sensors use negative feedback such that any changes in local field at the sensor are compensated by on-board-sensor electromagnetic coils. However, closed-loop operation is complicated since magnetic fields oriented in all three directions relative to the sensor affect the linearity of the response (Schofield et al., 2023; Tierney et al., 2019), meaning closed-loop operation is required on three independent axes.

These limitations mean that OPM-MEG systems are not yet the “final product” and there remains significant scope for development. Here, we demonstrate a new OPM-MEG platform with a miniaturised electronic control system that solves many of the practical limitations associated with the current generation of instrumentation. This system also enables three-axis closed-loop operation, allowing sensors to continue generating faithful measures of magnetic field, even in the presence of large background fields. In what follows, we report a study demonstrating the equivalence of our new system to established OPM-MEG hardware. Following this, we use an electromagnetic phantom (a device that makes “brain-like” magnetic fields) to confirm that closed-loop operates as intended. We exploit closed-loop operation to make measurements of brain activity as participants move freely – including a sitting-to-standing task. We exploit the miniaturised nature of the electronics by transporting our system between two laboratories and acquiring data at both sites. Finally, we pair our new system with EEG, demonstrating that we can acquire simultaneous OPM-MEG and EEG data.

## 2. MATERIALS AND METHODS

### 2.1 OPM-MEG SYSTEM COMPARISON

#### OPM-MEG systems and data acquisition

We initially aimed to compare two different OPM-MEG systems. Both comprised 64 triaxial OPM sensors (QuSpin Inc. Colorado, USA) each capable of measuring magnetic field in three orthogonal orientations, enabling data collections across 192 independent channels. The sensor design has been well documented (Boto et al., 2022; V. Shah et al., 2020) and will not be repeated in detail here; briefly, each sensor head is a self-contained unit incorporating a ^87^Rb vapour cell, a laser for optical pumping, on-board electromagnetic coils for field control within the cell and two photodiodes for signal readout. A beam splitter splits the laser output and associated optics projects two orthogonal beams through the cell, to enable triaxial field measurement. The median noise floor of the sensors was expected to be ∼15 fT/sqrt(Hz) in the 3-100 Hz range. This is marginally higher than the noise floor of typical single or dual axis OPMs due to the requirement to split the laser beam for triaxial measurement (Boto et al., 2022). Sensors from both systems were mounted in identical 3D-printed helmets (Cerca Magnetics Limited, Nottingham, UK) ensuring that the array geometry was the same for all measurements (see Figure 1A – inset photo). The arrays were housed in a magnetically shielded room (MSR) comprising four mu-metal layers and one copper layer to attenuate DC/low frequency and high frequency magnetic interference fields, respectively (Magnetic Shields Limited, Kent, UK). The MSR walls were equipped with degaussing coils to reduce remnant magnetisation prior to a scan. The MSR was also equipped with a matrix coil (Holmes et al., 2023) and a fingerprint coil (Holmes et al., 2019) – both capable of active field control (Cerca Magnetics Limited, Nottingham, UK). A single “acquisition” computer was used for OPM-MEG control and data acquisition; the paradigm (along with associated temporal markers (“triggers”) delineating the time at which stimuli were presented to the subject) was controlled by a second “stimulus” computer. Visual stimuli were presented via projection through a waveguide onto a back projection screen positioned ∼100 cm in front of the subject. We used an Optoma HD39 Darbee projector with a refresh rate of 120 Hz. Schematics of both systems are shown in Figure 1C.

**Figure 1:**
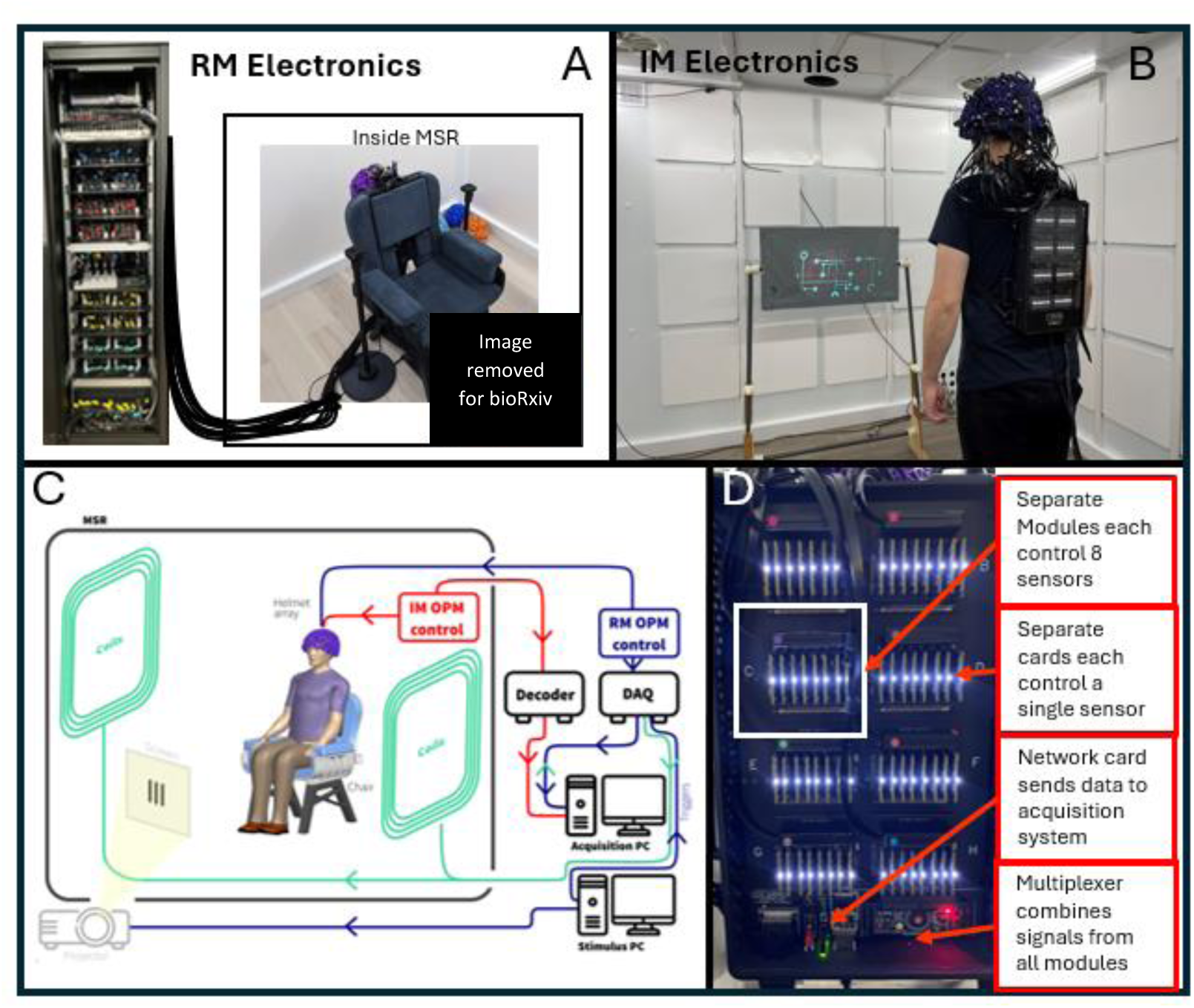
OPM-MEG systems. A) Rack mounted (RM) OPM-MEG system; sensor heads controlled via an electronics rack outside the MSR. B) Integrated miniaturised (IM) OPM-MEG system; all control and acquisition electronics contained on within a backpack worn by the subject. C) System schematic – valid for both systems with the major difference being the OPM electronics: Red pathways shows the IM system, blue the RM system. D) Photograph of the electronics for the integrated miniatured system.

The two OPM-MEG systems differed substantially in terms of their electronics control.

- **Established “Rack mounted” (RM) System:** In the case of our established system (Figure 1A) each sensor head was connected by a lightweight (2.2 gm^-1^) ribbon cable, 90 cm in length, to a backpack. The backpack housed 64 junction boxes in which the ribbon cables were connected to more robust cables (40 gm^-1^; 550 cm in length). 64 of these larger cables pass through waveguides in the MSR wall and connect to an electronics control rack positioned outside the MSR. Each sensor head is controlled independently by an electronics unit which provides all control signals (including temperature modulation (via proportional-integrative-derivative control) for the vapour cell and laser as well as on-board sensor coil control (including the modulation signals required for directional sensitivity (Cohen-Tannoudji et al., 1970)). The output of each sensor electronics unit includes three analogue signals which represent the three orthogonal fields measured by the sensor. These 192 separate signals are fed into a single data acquisition system (DAQ – sbRIO9637, National Instruments,) where they are digitised synchronously and sent to the acquisition computer. The DAQ also has 16 additional channels for digital and/or analogue triggers as well as peripheral analogue signals (here we only needed 6 triggers for the paradigm). The independently controlled sensors are synchronised via a 923 Hz sinusoidal modulation signal, and an externally generated 307 kHz sinusoid heater signal fed into each unit. The rack is powered by two 300 W power supplies. The rack is 2.02 x 0.6 x 0.6 m^3^ and weighs approximately 240 kg; the total weight of cabling between the backpack and the electronics rack is 14 kg.
- **New “Integrated Miniaturised” (IM) system:** For our new system all control electronics are housed in a backpack which is worn by the subject (Figure 1B) (*Neuro-1-electronics*, QuSpin, Colorado, USA – https://quspin.com/neuro-1-an-integrated-sensor-system-for-opm-meg/). Sensor heads were connected to the backpack by a ribbon cable; 2.2 gm^-1^ and 90 cm in length. Sensors are grouped into modules, with a single module controlling up to 8 sensors; each sensor is controlled by a single electronics card which – like the electronics units in the RM system – provides control signals for all on-board sensor components. Unlike the RM system, this device is capable of three-axis closed-loop operation in which the magnetic field is measured at each OPM along all three orthogonal axes, with the electronics effecting a negative feedback loop, whereby currents are applied to the on-board sensor coils to maintain zero field at the vapour cell. This linearises the OPM response to external fields and theoretically extends the dynamic range. Each electronics card contains its own DAQ and the outputs from all modules are digital. Those outputs are sent to a multiplexer and then to a network card. The backpack is connected via ethernet to a decoder positioned outside the MSR. This integrates the MEG signals with triggers sent from the acquisition PC, samples all signals synchronously, and passes them to the acquisition PC via an ethernet connection. (Note: the system contains 3 BNC inputs and a parallel port for digital triggers and 8 analogue-to-digital converters for peripheral signals, though again only 6 channels were required.) The only other connection to the backpack is a power cable. The backpack is 0.36 x 0.2 x 0.06 m^3^ and weighs approximately 1.8 kg.

Data were recorded from our RM system at 1200 Hz, and from our IM system at 375 Hz (this was the same for all experiments, except the ‘Sitting-to-standing task’ which was sampled at 1500 Hz). In both systems, data are stored on the same acquisition PC. For all recordings, participants were seated comfortably on a patient support located in the centre of the MSR. Prior to the recording, the inner walls of the MSR were degaussed, and the fingerprint coils energised using pre-determined currents. This ensured that the coils generated magnetic fields in the region surrounding the subject’s head that were equal and opposite to the fields typically observed in the MSR (these had been determined based on field measurements made across multiple previous experimental sessions (Rhodes et al., 2023)). The field surrounding the participant’s head was consequently < 500 pT. Participants were free to move throughout the recording. For our IM system, data for this experiment were recorded in *open-loop* mode.

We used optical scanning to determine how the helmet was positioned on the subject’s head. Immediately following MEG data acquisition, a 3D digitisation of the participant’s head (with the helmet in place) was acquired using a 3D structured light scan (Einscan H, SHINING 3D, Hangzhou, China). The 3D surface of the subject’s face was extracted from this scan and matched to the equivalent surface taken from a T1-weighted anatomical magnetic resonance image (MRI). This enabled a co-registration of the helmet to brain anatomy (Hill et al., 2020; Zetter et al., 2019). Following this, detail of the precise locations of the sensors within the helmet (generated by the 3D printing of the helmet itself) was added to give a complete description of sensor locations/orientations relative to anatomy.

#### Experimental paradigm

Our first aim was to demonstrate that our new system had similar performance to an established OPM-MEG system (Rea et al., 2022; Rhodes et al., 2023; Rier et al., 2023, 2024). To this end, we scanned the same individuals using both systems (see Figures 1A/B) and compared results. Two healthy participants (both male, ages 28 and 43) took part in the study, which was approved by the University of Nottingham Faculty of Health Sciences Research Ethics Committee. Each participant was scanned 6 times in each system over a period of 3 days (the order of scanning was counterbalanced). The scanning session was repeated at the same time each day for each participant (one participant session in the morning and one in the afternoon). The experiment consisted of a visuo-motor paradigm (Hill et al., 2022), containing 3 trial types:

1. **Circles trials:** A visual stimulus (a central, inwardly moving, maximum-contrast circular grating) was presented. The grating subtended an angle of 6° and was displayed for a duration of 1 s. This was followed by a (jittered) baseline period lasting 1.25 ± 0.2 s where a central cross was displayed. There were 60 circles trials per experiment. This stimulus is known to induce gamma oscillations in visual cortex (Hill et al., 2022; Hoogenboom et al., 2006; Iivanainen et al., 2020).
2. **Faces trials:** Visual stimulation was a photograph of a face, displayed on screen for a duration of 0.5 s, followed by a jittered rest period of duration 1.25 ± 0.2 s (during which a fixation cross was shown). A total of 120 faces trials was used. This task generates evoked responses in primary visual and fusiform areas (Hill et al., 2022).
3. **Catch trials:** Here, a cartoon character was displayed for 0.8 s. A total of 25 catch trials were presented and upon presentation, the subjects were asked to press a button with the index finger of their right hand. Such movements elicit robust modulation of beta oscillations in primary sensorimotor cortices (Pfurtscheller & Lopes da Silva, 1999).

The trial types were pseudo-randomised across the experiment, and the total experimental duration was 396 s. Prior to all sessions, empty room noise data were also collected using both systems.

#### Data analysis

Channel noise spectral densities were inspected visually and any channels that were either excessively noisy or had close to zero amplitude (i.e., not functioning) were removed. A trial-by-trial analysis was carried out and any trials containing excess artefact were also removed. Notch filters at the powerline frequency (50 Hz) and two of its harmonics, as well as a 1-150Hz band pass filter were applied. Finally, homogeneous field correction (HFC) was applied to reduce interference caused by distant sources (Tierney, et al., 2021). We used a beamformer spatial filter (Robinson & Vrba, 1998) to construct either pseudo-T or pseudo-Z statistical images showing the spatial signature of task induced change in source power or amplitude. We also used a beamformer to construct timecourses of electrophysiological activity at locations of interest (derived from locations informed by the statistical images – termed a ‘virtual electrode’). Specifically:

- **For Circles trials**, data were segmented to 0 to 2 s windows (relative to the onset of the circle) and filtered to the 52 to 65 Hz frequency band (as the circular grating is known to elicit a narrow-band response at 60Hz (Hoogenboom et al., 2006; Iivanainen et al., 2020)). A covariance matrix and beamformer weights were constructed using filtered data from all Circles trials; covariance matrices were regularised using a regularisation parameter equal to 1% of the maximum eigenvalue of the unregularized matrix (this is the case for all beamformer images derived in this paper). To make the pseudo-T-statistical image, we contrasted power in the 0 to 0.6 s (active) window to power in the 1.1 to 1.7 s (control) window, deriving a pseudo-T-statistic for voxels on a regular 4-mm grid covering the brain. To generate a virtual electrode, beamformed data local to the peak in the pseudo-T-statistical image were Hilbert transformed and the analytic signal derived. The absolute value of this signal was then used to give the envelope of oscillatory amplitude (*Hilbert envelope*) in the gamma band, which was trial averaged.
- **For Faces trials**, data were segmented to 0 to 1.5 s windows (relative to the presentation of a face) and filtered to the 2 to 40 Hz band. Covariance and weights were constructed using data from all Faces trials. To compute the evoked response, we first used a beamformer to reconstruct a virtual electrode at an anatomically defined point in the fusiform cortex (selected according to the automated anatomical labelling atlas (Gong et al., 2009; Hillebrand et al., 2016; Tzourio-Mazoyer et al., 2002)). Beamformed timecourses were averaged across trials, giving the evoked response. For the peak in the evoked response at ∼170 ms (i.e., a single point in time) we generated a pseudo-Z-statistical image to assess spatial signature for each experimental run.
- **For Catch trials**, data were segmented to −0.3 to 1.7 s windows (relative to the button press) and filtered to the beta (13 to 30 Hz) band. Covariance and weights were constructed using data from all Catch trials and the pseudo-T-statistical image contrasted power in the −0.3 to 0.3 s window to power in the 0.8 to 1.4 s window. A virtual electrode was generated for a location derived from the pseudo-T statistical image, using the Hilbert envelope to show the time evolution beta band oscillatory amplitude.

In all cases, data from both systems and all experiments were processed in the same way and results compared.

### 2.2 CLOSED-LOOP OPERATION

Our second aim was to demonstrate that closed-loop sensors enabled operation of our IM system in the presence of large background magnetic fields. For this we used a two-step approach, first employing a phantom to make known magnetic field signals, and then moving to a more naturalistic experiment in a human participant.

#### Phantom study

We used a dry-type current dipole phantom (Holmes et al., 2023; Oyama et al., 2015; Rier et al., 2023) to generate magnetic fields mimicking brain activity (henceforth called the *phantom field*). The phantom comprised a triangular electromagnetic coil (isosceles, 5-mm base and 45-mm height; made from a single turn of 0.56-mm diameter enamelled copper wire. The ends of the wire were twisted to avoid any stray magnetic fields.) The phantom was enclosed in a Perspex cylinder and glued to an empty OPM sensor casing allowing it to be fitted inside the OPM helmet with a known position and orientation relative to the sensor array. We performed two experiments to investigate the capabilities of closed-loop sensor operation.

- ***Response linearity in zero background field***: We first tested whether closed-loop operation had any effect on measurements being made in ‘zero’ background field (practically this means a background field at each OPM of <500 pT; well within normal (open loop) operational range). A sinusoidal current of frequency 17 Hz was applied to the phantom for 2 s followed by 1 s at zero current, to mimic experimental trials. The amplitude of the current waveform was varied between 8 values [0.01, 0.02, 0.05, 0.08, 0.1, 0.2, 0.5, 0.8, 1] mA, corresponding to an expected magnetic field between 20 and 200 pT at the OPM with the largest response. This process was repeated 8 times, and a trigger signal was used to mark periods in the data when the phantom was active. The experiment took ∼7 minutes and was repeated twice, once with sensors in open-loop mode, and a second time with sensors running in closed-loop mode. To check the linearity, we took the sensor outputs (for all channels) for both datasets, segmented data into 2-s periods where the phantom was active using the trigger signal, and applied a Fourier transform to compute a magnitude spectrum. We extracted the magnitude of the 17-Hz phantom signal for each channel, for all dipole amplitudes and trials (i.e., 64 measurements per channel per condition) and plotted all the open-loop sensor amplitudes against the same measures made using closed-loop operation. We expected that in this (zero-background-field) case, the values would be approximately the same with a linear relationship, meaning that closed-loop operation is not affecting measurement accuracy.
- ***Sensor response in non-zero background fields***: In the absence of closed-loop sensor operation, we would expect the presence of a large background field to affect the OPM response according to the solution to the Bloch equations which govern the polarisation of the rubidium gas (see Cohen-Tannoudji et al. (1970) for a complete description) – in general, as background field (in any orientation relative to the sensors) is increased, one would expect the individual OPM-response to the phantom field to decrease in amplitude. To investigate this, we used the matrix coil (Holmes et al, 2023), embedded in the walls of the MSR, to generate a controlled *background field*; this was uniform across the volume occupied by the OPM helmet and oriented vertically. We also generated a phantom field using the same current waveform as before, but only at a single amplitude – 1 mA – which produced the largest fields. We applied the waveform 5 times in zero applied background field, then stepped the background field up from 0 nT to 8 nT in 81 steps of 0.1 nT, repeating the measurement of phantom field waveforms after each step up in background field. The whole experiment took around 20 minutes and was repeated twice with sensors in open- and closed-loop mode. We segmented the data and computed the magnitude spectrum as before, again plotting corresponding open-loop versus closed-loop measurements of the phantom field, for all trials and all values of background field (405 measurements per channel per condition). Here we expected that closed-loop operation would result in faithful reconstruction of the expected field, whereas with open-loop operation, the measured field would diminish with increasing background field, and, unlike the previous experiment, there would be no linear relationship between the two operational modes.

#### Sitting-to-standing task

As a further demonstration of closed-loop operation, we recorded data during a naturalistic task. A single participant (male, aged 28) took part in the study. In our ‘sitting-to-standing’ task, trials lasted 8 s and were cued by two alternating auditory stimuli (either a 1400-Hz tone or a 1000-Hz tone, both lasting 1 s). On hearing the 1400-Hz cue, the participant moved from sitting to standing; on the 1000-Hz cue, the participant went from standing to sitting. Whilst moving they performed abductions of their right index finger. As the head moves it will experience a changing magnetic field; we purposely did not degauss the MSR or engage active field compensation prior to the measurement, to maximise the field change that would be experienced by the sensors. (We expected that the field change would be of order 2-3 nT – sufficient to take OPMs operating in open-loop outside their dynamic range.) With closed-loop operation, we hypothesised that the beta band modulation induced by the movements would be successfully recorded, despite the large shift in background field.

Data were processed using a similar pipeline to that described above: Following bad channel/trial rejection, data were segmented to 0 to 8 s windows (relative to the auditory cues) and filtered to the 13 Hz to 30 Hz band. Covariance and beamformer weights were constructed using data from all trials and a pseudo-T-statistical image contrasted oscillatory power in the 2 to 3 s window to that in the 6.5 to 7.5 s window. To examine signal dynamics, we constructed a time-frequency spectrogram (TFS). The pre-processed data were frequency filtered into the 1-150 Hz band and data covariance and beamformer weights estimated. A virtual electrode was generated for a location derived from the peak in the pseudo-T statistical image. The resulting (broadband) beamformed data were filtered into a set of overlapping frequency bands, and the Hilbert envelope computed for each band; this was averaged across trials and concatenated in frequency. Baseline correction was applied by subtracting the time average of oscillatory amplitude, for all bands, in the control window. The spectrum was normalised by oscillatory power in the control window. The result was a TFS showing relative change in oscillatory amplitude, for all frequencies, over time.

### 2.3 DEMONSTRATING PORTABILITY

One advantage of our IM system is that the small and lightweight electronics makes it transportable. Our third aim was to provide a proof-of-principle of this portability, by moving the system between two laboratories and recording data in a group of healthy volunteers. The two sites differed as follows:

- **Site 1: University of Nottingham, Nottingham, UK**: The system was housed in a MSR (Magnetic Shields Limited) with internal dimensions 3 x 3 x 2.4 m^3^, and walls comprising four layers of mu-metal and a single copper layer. The room was equipped with degaussing, and both matrix and fingerprint coils for active field control. All experiments were approved by the University of Nottingham Faculty of Health Science ethics committee.
- **Site 2: Hôpital Erasme, Bruxelles, Belgium:** The system was housed in a compact MSR (Magnetic Shields Limited) with internal dimensions 1.3 x 1.3 x 2.4 m^3^, and walls comprising two layers of mu-metal and a single layer of copper. The room is equipped with degaussing coils and a “window coil” for active field control (Holmes et al., 2022). Hôpital Erasme’s Ethics Committee approved this study (P2019/426) and participants gave written informed consent.

The same IM system (in closed-loop mode) was used in both laboratories, and was transported between sites in two suitcases, via rail. Five individuals took part in the experiment at each site (2 female and 3 male aged 25-33 at site 1 and 1 female, 4 male aged 27-47 at site 2).

The experimental paradigm was a motor task; a single trial comprised 3 s of a right index finger abduction, followed by 3 s rest. A total of 100 trials were collected over 2 runs of 50 trials. All data were collected at a sampling rate of 375 Hz. Data preprocessing was as described above, except a template brain warping method (Rier et al., 2024) was used to create a ‘pseudo-MRI’ for each subject as opposed to an anatomical MRI. Pseudo-T-statistical beamformer images were derived for the beta band (13 Hz to 30 Hz), by contrasting active (0.5 s to 1.5 s) and rest (3.5 s to 4.5 s) windows. TFSs were derived from locations of interest derived from peaks in the images.

### 2.4 CONCURRENT OPM-MEG EEG

EEG remains the most common clinical metric of brain function, and has proven utility in disorders including epilepsy, dementia, head injury, sleep disorders and encephalitis (Ding & Yuan, 2013). Previous work suggests that OPM-MEG can outperform EEG in terms of sensitivity, spatial resolution, and practicality (Boto et al., 2019). However, in practice it would be advantageous to acquire data from both modalities simultaneously; this would not only enable integration of signals (which has advantages (Aydin et al., 2015; Feys, et al., 2023; Yoshinaga et al., 2002)) but also allow clinicians to benefit from the improved performance of OPM-MEG whilst still having access to EEG data (Feys, et al., 2023). The use of simultaneous scalp EEG during MEG recordings in epileptic patients is mandatory according to international clinical practice guidelines (Bagić et al., 2011, 2023; Feys, et al., 2023). It allows for a better classification of physiological vs. pathological brain activities (Rampp et al., 2020). It also provides a truly complementary measure to MEG, such that neural sources “silent” in one modality (EEG vs. MEG) can be detected in the other modality (Mosher & Funke, 2020). The simultaneous collection of OPM-MEG and EEG data has been demonstrated previously (Boto et al., 2019; Feys, et al., 2023) – here our aim was to show that it was also possible using our IM system.

To this end, we employed a 64-channel MEG-compatible EEG system (Brain Products GmbH, Munich, Germany) comprising an EEG-cap (with passive, MEG compatible, AgCl electrodes), signal amplifiers, a power pack, and a data acquisition laptop. During data acquisition, the ground electrode was AFz and the reference electrode was FCz. 63 channels were attached to the scalp and one to the subject’s back to measure the electrocardiogram (ECG). Conductive gel (Abralyte 2000) was used to connect electrodes to the head with all impedances kept below 10 kΩ. Our 192-channel OPM-MEG helmet was placed over the top of the EEG cap and was otherwise operated (in closed-loop mode) as described above. Triggers from the stimulus PC were split so that simultaneous markers appeared in the OPM-MEG and EEG data, allowing data alignment. Five subjects took part in the experiment (1 female and 4 male, age 27-33). A single recording comprised 60 Circles trials and 80 Faces trials – though here we only analysed the circles trials to observe induced beta and gamma effects. In this experiment, participants were asked to complete a finger abduction with their right finger during the Circles trials. The experiment was carried out twice per subject, once when the participant was asked to remain still and once when they were asked to make natural head movements (head movements were tracked via infra-red markers placed on the helmet, tracked via OptiTrack (Natural Point Inc.) motion tracking cameras. Two 3D digitisations were taken: one of the participant wearing the OPM helmet (so that co-registration could be carried out, as previously described) and one of the EEG cap. Electrode positions were found from the digitisation and matched to a layout of the cap by first manually point-matching specific electrodes and then using an iterative closest point (ICP) algorithm. The electrode array was then co-registered to the MRI by matching fiducial points on the digitisation and MRI.

OPM-MEG processing was as described above. EEG data preprocessing comprised removal of bad channels and trials via visual inspection. Following this, data from remaining circles trials were segmented, concatenated, and filtered (independently) to both the beta (13-30 Hz) and gamma (30-80 Hz) bands. We constructed an EEG forward model using a 3-shell boundary element model implemented in Fieldtrip (Oostenveld et al., 2011). We then processed the data from both EEG and MEG using the same beamformer approach as was used in OPM-MEG (forward model and beamformer code available from https://github.com/SCColeman/EEG_beamformer). Covariance matrices were generated using band limited data. Pseudo-T-statistical images were derived on a 4-mm grid spanning the whole brain volume. For visual gamma effects, we contrasted the 0.2 s to 0.8 s active window with the 1.2 s to 1.8 s control window; to look for sensorimotor beta modulation, we contrasted a 0.5 s to 1.0 s active window with a 1.5 s to 2.0 s control window. For the beta modulation, we derived TFSs from the peak location found independently in both EEG and OPM-MEG. For the visual gamma effects in both the EEG and OPM-MEG, TFSs were derived from an anatomically defined location in the visual cortex (selected by taking the centre of mass of the calcarine region according to the automated anatomical labelling atlas (Gong et al., 2009; Hillebrand et al., 2016; Tzourio-Mazoyer et al., 2002)). This was due to inconsistency in the localisation of the gamma response in EEG. This process was repeated in OPM-MEG for consistency.

## 3. RESULTS

### 3.1 NOTES ON SYSTEM OPERATION

Our IM system had a representative noise floor of 14.0 fT/sqrt(Hz) (median across sensors in the 3-100 Hz band). This was compared to 12.3 fT/sqrt(Hz) for our RM system. We acquired data in 42 experiments using our IM system (2 subjects x 6 comparison scans; 4 phantom experiments; 1 sitting-to-standing task; 2 sites x 5 subjects x 2 runs cross site study, and 5 OPM-MEG/EEG experiments). Across all 42 experiments, we lost on average 3 ± 5 channels (mean ± standard deviation). These lost channels were typically due to an OPM sensor head becoming detached from the ribbon cable. The system typically took ∼60 s to start the 64 sensors (this procedure includes heating all sensors, locking laser frequencies (and temperatures) and optimising control parameters). Zeroing the magnetic field (using on-board-sensor coils) and calibrating each sensor took a further ∼60 s. The total system set up time (including degaussing and field nulling) was approximately 3 minutes. This was a similar set up time to the RM system.

### 3.2 OPM-MEG SYSTEM COMPARISON

Figure 2 shows the results of the comparison between our RM and IM systems. Results for a single subject are shown (averaged across all 6 runs); an equivalent figure for the second subject is provided in Supplementary Material. Panel A shows beta modulation during the button press. In both systems, the largest beta modulation was localised to the left primary sensorimotor cortices (due to movement of the right index finger) and the timecourse shows a clear movement induced reduction in beta amplitude, as expected. Figure 2B shows gamma modulation during presentation of the Circles stimuli. Here, the largest stimulus induced increases were in primary visual areas and the expected increase in gamma amplitude during stimulus presentation is observed. Figure 2C shows evoked responses to face presentation. The images show the spatial signature of the evoked response measured at a latency of ∼170 ms, which was predominantly in the fusiform area.

**Figure 2:**
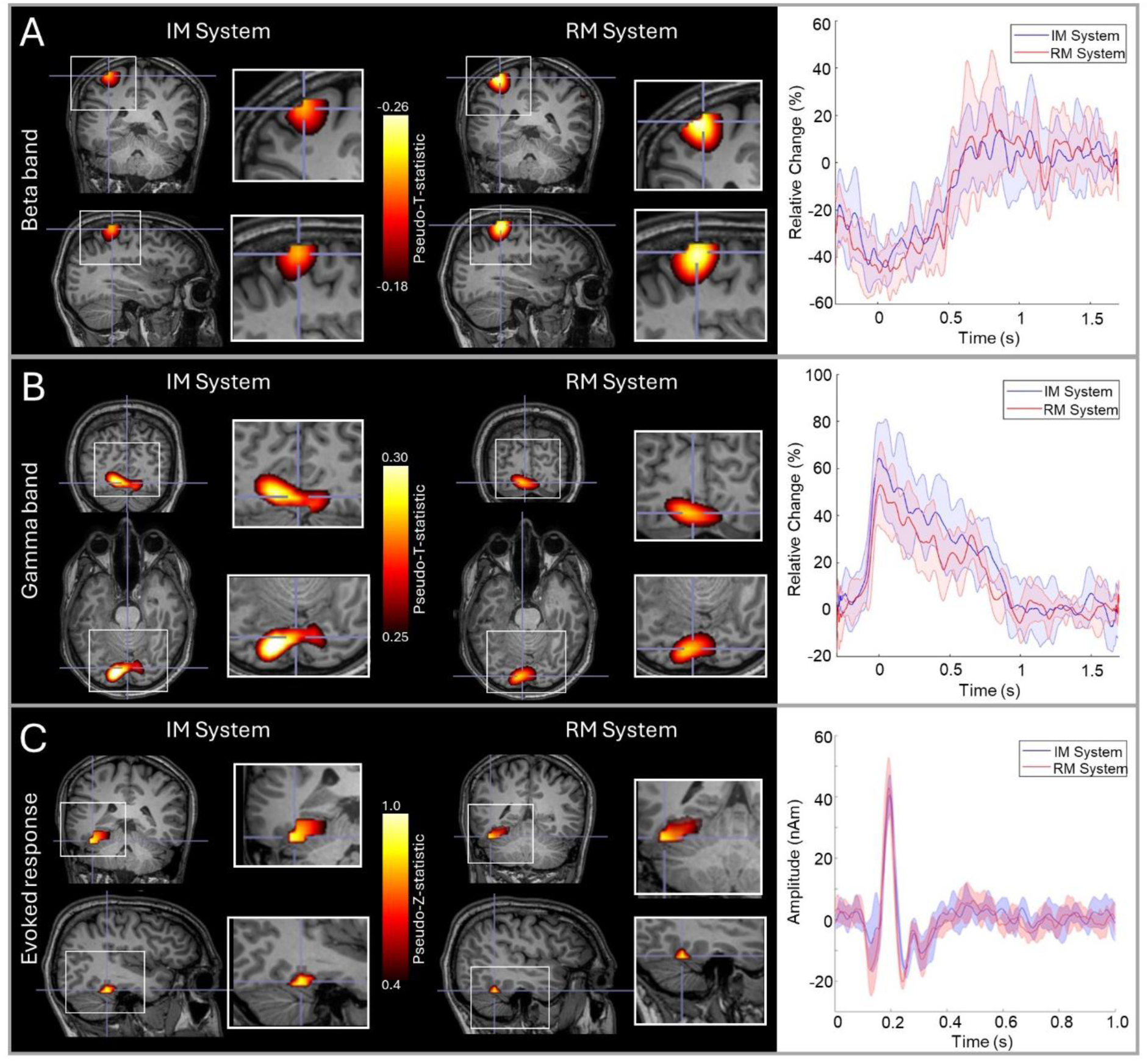
RM and IM system comparison. A) Beta band responses to a finger movement; in the images on the left the overlay shows the location of maximum beta modulation and the timecourses on the right show the time evolution of beta band amplitude. B) Gamma responses to visual stimulation; images show the locations of gamma modulation and timecourses show evolution of gamma band amplitude. C) Evoked responses to face presentation; images show the location of the highest evoked power and timecourses show trial averaged evoked responses. In all three cases, data are averaged across 6 runs; images from both systems are shown and in the timecourse plots, red represents the RM system, blue the IM system and the shaded areas represent standard deviation over runs. Data are shown in a single subject (S2). Data from the second subject are shown in Supplementary Material.

To quantify the spatial agreement between runs, we measured the Euclidean distance between the voxels showing the largest modulation for all three responses (beta, gamma, and evoked). This was done in three ways:

1. **Individual runs – between system:** In a single participant, for each measurement (beta, gamma and evoked response) we had 6 pseudo-T/Z-statistical images for each system. This gives 36 possible comparisons between systems (run1 – to run 1; 1 – 2; 2 – 2 etc.) For each, we found the Euclidean distance between peak voxels. We then found the mean and standard deviation of these values.
2. **Individual runs within system:** From the 6 runs in a single system there are 15 possible comparisons (i.e., runs 1-to-2; 1-to-3; 2-to-3 etc). For each within system comparison, we again measured Euclidean distance between the peak voxels, computing the mean and standard deviation.
3. **Averaged runs:** We measured the Euclidean distance between the peaks in the pseudo-T/Z-statistical images averaged across runs in the same system.

The results of these three analyses, which were computed for each subject, are shown in Table 1. The spatial discrepancies between runs tended to be larger for the gamma and evoked experiments than for the beta experiments. However, in all cases, the within system comparison is not significantly different to the between system comparison, suggesting that there is no major difference in spatial localisation between systems.

**Table 1:** Spatial robustness: All values represent Euclidean distances in mm between peaks in pseudo-T-statistical images. Comparisons are made between systems for individual runs, within systems for individual runs, and between systems for the averages across runs, for two subjects (S1 and S2).

| Distance | Individual Runs (mm) |  |  |  |  |  | Averages (mm) |  |
| --- | --- | --- | --- | --- | --- | --- | --- | --- |
|  | RM to IM |  | Within RM |  | Within IM |  | RM to IM |  |
| Response | S1 | S2 | S1 | S2 | S1 | S2 | S1 | S2 |
| Beta | 11.1 ± 7.7 | 8.0 ± 4.0 | 13.0 ± 9.1 | 8.8 ± 2.8 | 8.0 ± 4.1 | 8.7 ± 5.0 | 6.8 | 1.6 |
| Gamma | 34.0 ± 13.8 | 17.0 ± 7.8 | 31.1 ± 22.7 | 20.4 ± 7.6 | 30.0 ± 20.2 | 16.6 ± 8.3 | 14.2 | 2.7 |
| Evoked | 19.9 ± 12.8 | 21.7 ± 14.5 | 16.6 ± 11.1 | 16.4 ± 11.1 | 22.7 ± 12.0 | 29.5 ± 16.7 | 11.0 | 6.7 |

To quantify the temporal agreement between runs we used Pearson correlation between the reconstructed timecourses of either induced or evoked activity. Again, we used three measures:

1. **Individual runs – between system:** We computed all 36 possible values of correlation between all pairs of experiments in the RM and IM system.
2. **Individual runs within system:** We calculated 15 correlation coefficients representing the similarity of experimental timecourses within each system (i.e., 30 measures in total).
3. **Averaged runs:** We calculated the correlation between timecourses averaged across runs.

As before, these calculations were carried out for each subject and paradigm separately. Results are shown in Table 2. First, note that all values of correlation are high (on average 0.77 for individual runs and 0.91 for averaged runs) indicating that both systems are reliable. Again, we found no measurable difference between the within-system correlations (mean = 0.76) and between-system correlations (mean = 0.77).

**Table 2:** Temporal robustness for subject 1 (S1) and subject 2 (S2): All values represent Pearson correlation between trial averaged timecourses of either beta modulation, gamma modulation, or evoked responses, for two subjects (S1 and S2).

| Correlation | Individual Runs |  |  |  |  |  | Averages |  |
| --- | --- | --- | --- | --- | --- | --- | --- | --- |
|  | RM to IM |  | Within RM |  | Within IM |  | RM to IM |  |
| Response | S1 | S2 | S1 | S2 | S1 | S2 | S1 | S2 |
| Beta | 0.81 ± 0.05 | 0.75 ± 0.06 | 0.81 ± 0.03 | 0.78 ± 0.05 | 0.80 ± 0.06 | 0.72 ± 0.07 | 0.96 | 0.96 |
| Gamma | 0.67 ± 0.08 | 0.81 ± 0.07 | 0.67 ± 0.08 | 0.79 ± 0.05 | 0.69 ± 0.08 | 0.85 ± 0.05 | 0.91 | 0.95 |
| Evoked | 0.65 ± 0.2 | 0.84 ± 0.06 | 0.62 ± 0.16 | 0.89 ± 0.04 | 0.84 ± 0.07 | 0.81 ± 0.06 | 0.75 | 0.91 |

### 3.3 CLOSED-LOOP OPERATION

Figure 3 shows the results of our phantom experiments. Panel A shows the amplitude of the phantom field (i.e., the amplitude of the 17 Hz component of the Fourier spectrum) measured using open-loop (plotted on the x-axis) versus closed-loop (plotted on the y-axis) operation. All data were acquired in zero-background field and the black line represents the line of equality. The linear relationship shows that using closed-loop mode makes no difference to the measurements. Panel B again shows open-loop measures of the phantom field plotted against closed-loop values. However, here the background field was allowed to vary between 0 nT and 8 nT (this is represented by the colour of the data points). As background field was increased, the fields measured in open-loop mode decrease as expected, but the fields measured in closed-loop remain the same.

**Figure 3:**
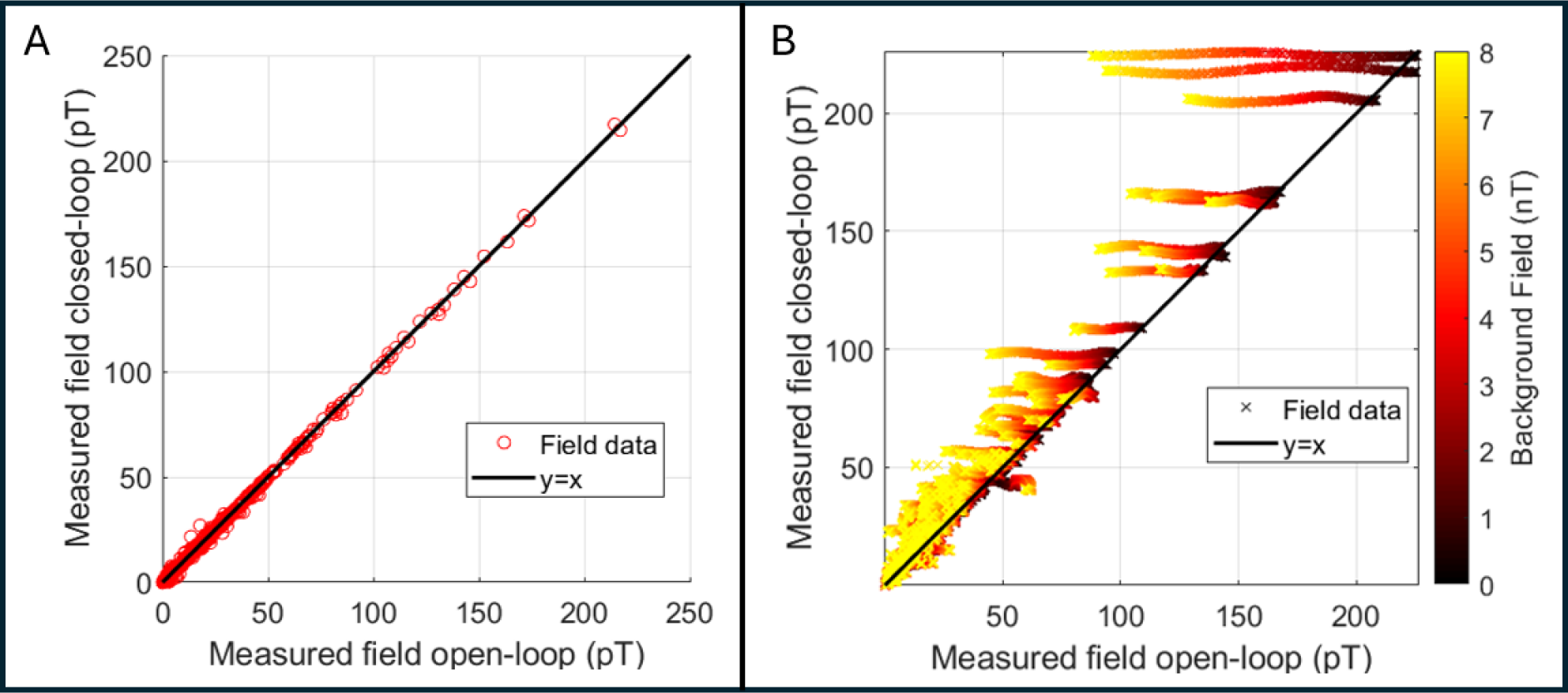
Phantom experiments to characterise closed-loop operation. A) Measured phantom field amplitudes with zero background field. Measurements in closed-loop mode plotted against equivalent measures in open-loop mode. Note the linear relationship showing that closed-loop operation makes no difference to field measurement in low background field. B) Phantom fields measured with a varying background. Again, measures acquired with closed-loop on are plotted against equivalent measures in open-loop mode. Here, the background field was allowed to vary between 0 and 8 nT (delineated by the colour of the points). As background field is increased, the open-loop fields decrease, yet the closed-loop fields remain at approximately the same value.

To make a quantitative assessment of these effects, we took all the measurements and subtracted the value measured with zero background field (i.e. we removed our best assessment of the true value) for the 10 sensors measuring the largest phantom field. We then computed the difference from zero as an estimate of the error introduced by the non-zero background. In closed-loop mode, this error was 0.5 ± 0.35% (mean ± standard deviation) whilst in open-loop mode it was 14.2 ± 14.9 %, for a range of background fields of 0 nT - 8 nT.

Figure 4 shows the results of our sitting-to-standing task. Figures 4A and C show pseudo-T-statistical images of beta modulation and a TFS extracted from the peak in primary sensorimotor cortex, respectively. The largest beta modulation was localised to the bilateral sensorimotor regions, extending from the hand area medially to the areas responsible for leg movement (recall that the task involved finger movement whilst standing up, so this is to be expected). The TFS showed clear beta band desynchronisation in the first 4 seconds of each trial whilst the subject was moving. Figure 4B shows the raw magnetic field data measured by the sensors. Most sensors show a background field shift, generated by movement, of >1.5 nT – this is more than the dynamic range of the sensors when running in open-loop mode. Despite these large field shifts, the sensors maintain operation. While these measurements would be possible with the sensors running in open-loop, the accuracy of the signals would be significantly impeded by both gain and CAPE errors (Borna et al., 2022).

**Figure 4:**
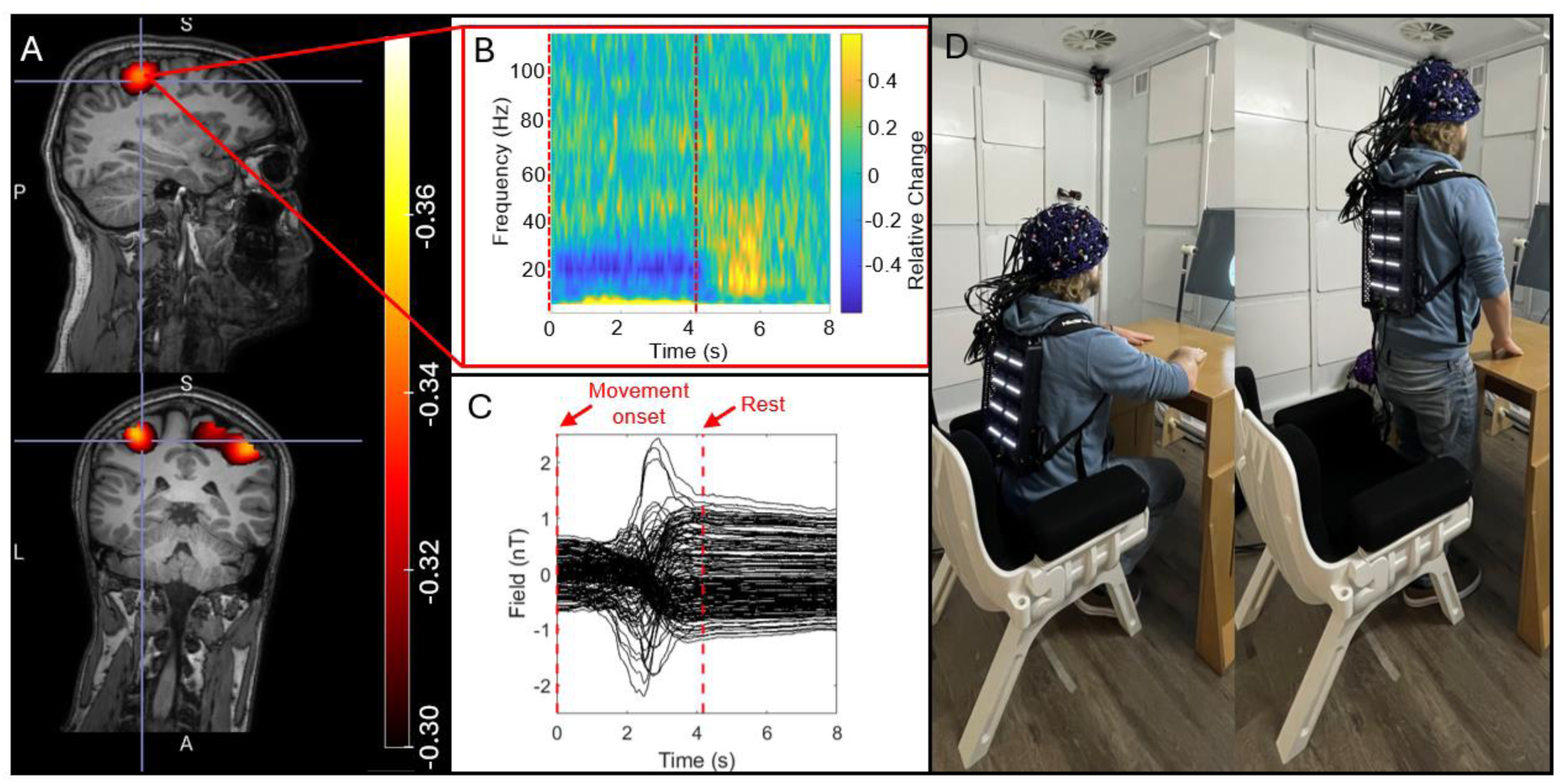
Sitting-to-standing task. A) The spatial signature of beta modulation induced by the task. B) The raw magnetic fields measured by the channels, showing that sensors travelled through a ∼2 nT background field as the participant moved from sitting to standing C) a TFS from sensorimotor cortex showing the time frequency evolution of neural oscillations. D) a re-enactment of the task to demonstrate range of motion.

### 3.4 CROSS SITE COMPARISON

Figure 5 shows the results of our cross-site study where the IM system was transported between laboratories in Brussels and Nottingham. Figure 5A shows pseudo-T-statistical images of beta modulation induced by finger movement. The upper panels show Brussels data and the lower panels show Nottingham data – all results are averaged across five subjects at each site. The TFSs in panel B show the time frequency dynamics at the image maxima. Peaks in the averaged images were separated by 1 mm and temporal correlation was 0._A_96. For direct comparison, Figure 5C shows the envelope of oscillatory amplitude in the beta band, averaged over trials and subjects. Note the high level of similarity.

**Figure 5:**
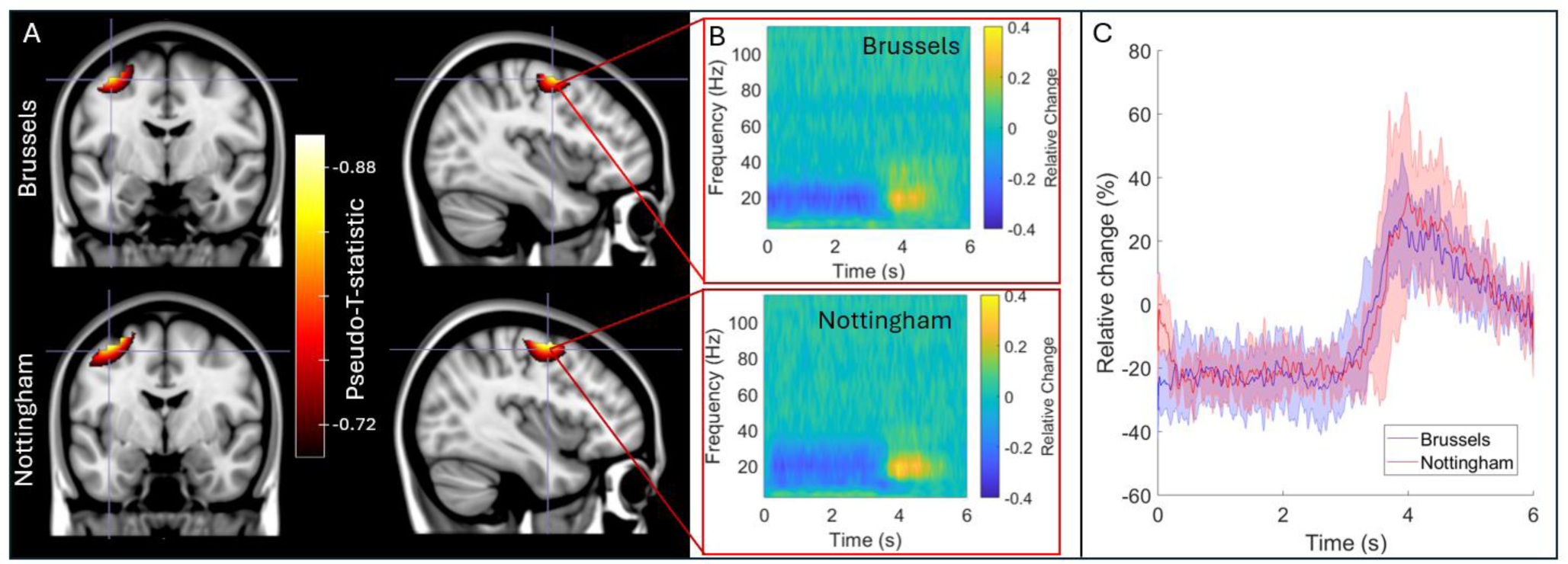
cross site comparison. A) Pseudo-T-statistical images for the group average beta band effects at the two sites. In both cases, the upper panels show data acquired at the Nottingham site and the lower panels show the Brussels site. B) TFSs extracted from the image peaks. C) Trial-averaged reconstructed timecourses filtered to the beta band, extracted from the image peaks.

### 3.4 CONCURRENT OPM-MEG/EEG

Finally, Figure 6 shows the results of our concurrent OPM-MEG/EEG experiments. Here five people took part in the experiments, however averages are shown in only four participants since the EEG recording failed in one. Figure 6B shows the extent of the natural movements carried out by subjects during the scans; bar charts show maximum translation (bottom) and rotation (top); the bars show the mean across subjects, and the individual data points show results for the four subjects. Figures 6C and D show concurrently acquired OPM-MEG and EEG data averaged across subjects. Panel C shows the beta band modulation during finger movement and panel D shows gamma modulation by the visual stimulus. In both cases, pseudo-T-statistical images and TFSs are included. As expected, EEG and OPM-MEG show similar effects; both allow recording of beta and gamma oscillations showing the viability of concurrent recordings. However, whereas localisations for OPM-MEG are as expected (primary motor and visual cortices for the beta and gamma effects respectively) the localisations do not appear as accurate for the EEG data – this will be discussed further below.

**Figure 6:**
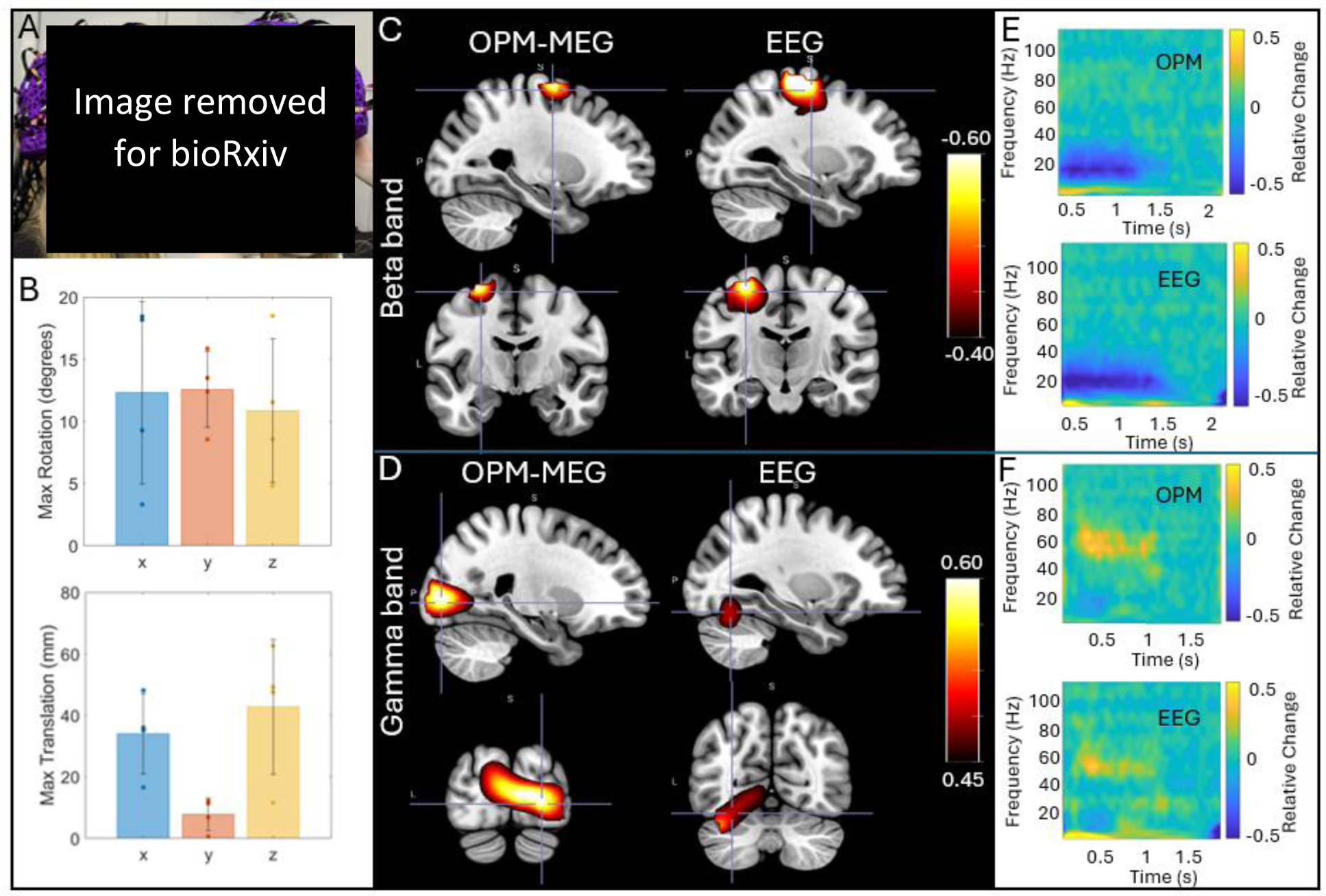
Concurrent OPM-MEG/EEG. A) A participant wearing an EEG cap and OPM-MEG helmet. B) Data were recorded during natural head movements: the maximum translations and rotations made by the subjects across the experiments are shown. Bars represent mean across subjects; data points show values for each individual subject. C) and D) show group average beta and gamma effects, respectively. In both cases pseudo-T-statistical images and associated TFSs (from the minima for beta and a central point in the visual cortex for gamma) in those images are shown for EEG and MEG. All data were recorded in the presence of movement. The static case is shown in Supplementary Material.

## DISCUSSION

Our overarching aim was to demonstrate a new OPM-MEG system with integrated and miniaturised electronics and test its viability for assessment of human electrophysiological function. Our primary demonstration saw the new IM system used multiple times in two subjects, to provide a comparison with an established OPM-MEG device, which has previously been extensively validated (Boto et al., 2022; Rea et al., 2022; Rier et al., 2023, 2024), including against conventional MEG (Boto et al., 2021; Hill et al., 2020; Rhodes et al., 2023). The results obtained with the two systems showed striking consistency; for beta band modulation in the motor task, the locations of maximum beta modulation were within ∼10 mm between systems. For gamma and evoked responses, we saw slightly lower consistency (∼20-30 mm) – this is likely due to larger source extent: in the case of gamma activity the centrally presented visual stimulus activates the visual cortex across both hemispheres, and the precise location of the peak modulation can appear on either side of the calcarine fissure, dependent on factors such as gaze position. Similarly, in the case of the evoked response, activity can be localised to both primary visual and fusiform regions. Most importantly, however, in beta, gamma, and evoked measurements the between system consistency did not differ significantly from the within system consistency – suggesting that systems are not measurably different. Source timecourses were highly reproducible across systems with an average correlation of ∼0.75 for individual runs, and >0.9 for averages of multiple runs in the same subjects. Overall, these results show that the two systems provide equivalent performance. Importantly, not only does this validate the miniaturised electronics, but also shows that the presence of this electronics inside the MSR (worn as a backpack) does not generate significant magnetic interference at the OPM sensors that cannot be rejected in post-processing through methods such as homogeneous field correction (Tierney, et al., 2021) and beamforming (Brookes et al., 2021).

The addition of closed-loop sensor operation is a significant technical milestone. The dynamic range of OPMs is low (±1.5 nT) and maintaining an environment where fields are limited to this range is challenging. Firstly, even inside magnetic shields, environmental changes in magnetic field over time (e.g. caused by local infrastructure) can be much larger than the dynamic range of the OPMs (e.g. (Hill et al., 2022)). Secondly, most MSR’s have a temporally static field ranging between 3 and ∼30 nT. These fields are typically nulled within each OPM using on-board sensor coils at the start of an experiment and so have no effect on measurements, if OPMs remain stationary. However, if the head is allowed to move in this static field, then the OPMs “see” a field that changes in time and can be taken outside their dynamic range.

The effects of both environmental field shifts and subject movement are further complicated by the orientation of the background field relative to the sensor: Let us assume we want to measure a field of interest, 𝐵_𝑥_, oriented in the x-direction. However, the measurement must be made in a background field described by the vector [𝐵_𝑏𝑥_, 𝐵_𝑏𝑦_, 𝐵_𝑏𝑧_]. Ideally, the sensor output along the x-direction would be 𝐵_𝑥_ + 𝐵_𝑏𝑥_ (i.e., the field of interest plus the background); indeed this is the case for low values of 𝐵_𝑏𝑥_, 𝐵_𝑏𝑦_ and 𝐵_𝑏𝑧_ (when the sensor is operating within its dynamic range). However, when background fields become large, then the presence of a background field along the measurement direction, 𝐵_𝑏𝑥_, will cause the OPM response to become non-linear, which manifests as a drop in sensor gain, meaning that the measurements of both 𝐵_𝑥_ and 𝐵_𝑏𝑥_ will be smaller than their true values. If 𝐵_𝑏𝑦_ and 𝐵_𝑏𝑧_ are large, this also causes a drop in OPM gain (again the measurement of 𝐵_𝑥_ diminishes); in addition, changes in 𝐵_𝑏𝑦_ and 𝐵_𝑏𝑧_ over time can manifest as a change in 𝐵_𝑥_ – this is known as cross-axis projection error (CAPE) (Borna et al., 2022). The ability to make closed-loop measurements, where measured fields at the vapour cell are compensated in real time by equal and opposite fields generated by the on-board-sensor coils offers a solution to this significant problem by making the sensor robust to changes in background field. However, because background fields in any orientation cause inaccuracies, complete 3-axis closed-loop operation is critical.

Here, we assessed closed-loop sensor operation in two ways. Firstly, we operated sensors in zero background field, in closed- and open-loop mode. The linear relationship shown in Figure 3A suggests that closed-loop operation (with zero background field) made no difference to the sensor output (within the 0-200 pT range of fields that we employed). Our second phantom experiment (Figure 3B) showed that, when background fields were applied, open-loop operation becomes inaccurate, with (in this particular experiment) sensor outputs dropping by ∼14% (on average over the 10 OPM channels measuring the largest phantom fields). However, when operating in closed-loop mode, the equivalent sensor error is within 0.5%. Importantly, the background field (which was oriented vertical relative to the MSR) would have intersected sensors with a number of different orientations since the sensors were positioned in a helmet, meaning that the triaxial nature of closed-loop operation was tested. The result therefore shows, for the first time, that 3-axis closed-loop operation can effectively linearise the response of an OPM in background fields of varying orientation up to and including 8 nT. In theory, the same approach can be used to reach even higher background field values (limited only by the range of field that can be generated by the on-board-sensor coils – ±50nT). This should be a topic of future system development.

One of the biggest advantages of OPMs over conventional MEG is that the sensors move with the head, enabling movement during a scan. This was shown here via two experiments – the sitting-to-standing task, where a participant transitioned from being seated to standing (Figure 4), and the concurrent EEG measurements, where participants made natural head movements. This type of investigation has significant utility – for example, the sitting-to-standing task is used widely used in clinical assessments of lower limb function, mobility, and fall risk across a range of conditions (e.g. (Nocera et al., 2013)). Similarly, children find it hard to keep still in conventional scanning environments, as do patients who may often be nervous, in pain, or even undergoing seizures (Feys, Corvilain, Van Hecke, et al., 2023; Hillebrand et al., 2023). The ability to scan people whilst moving therefore offers a more effective environment to gather MEG data. The introduction of our IM system offers two improvements in this area: Firstly, the new backpack-mounted electronics reduces the weight of cabling carried by the subject by a factor of 105. This makes naturalistic experiments where subjects are standing or walking much more practical, with only two cables to the backpack which can be easily managed (distinct from 64 cables for the RM system). Secondly, closed-loop operation means that sensor output remains linear even when the sensors move through a large background field; this enables sensors to keep working when the subject moves (as demonstrated in Figure 4). In previous demonstrations of free movement (Holmes et al., 2018, 2023; Rea et al., 2022), sensor linearity has been enabled by the use of large coils – either mounted on planes each side of the subject, or within the MSR walls – which create a zero field volume enclosing the head. In principle, closed-loop operation offers an alternative to use of such coils. However, it is important to note that larger coils not only linearise sensor output, but because they zero the field across the entire head volume, they also reduce artefacts caused by movement. Closed-loop operation linearises output but *doesn’t remove artefact*. This means that large scale coils are still a critical requirement for OPM-MEG instruments. However, it is worth noting that the increased accuracy in signal measurement from closed-loop operation may allow better characterisation and therefore removal of artefacts from data.

As part of the evaluation of our IM system, we exploited the compact and lightweight nature of the system by transporting it between two laboratories in different countries. Practically, this proved to be relatively straightforward – the system fitted into two suitcases and was easily transportable between sites. Such portability is somewhat tempered by the requirement for a MSR at both sites. Nevertheless, it offers new opportunities: For example, it becomes possible to design and build a single optimised array and scan individuals at multiple sites. This would be of significant utility – for example, to expand the numbers of patients that could be scanned in a single study by taking the same system to the patients. Equally, one can imagine shipping a system to more clinically oriented sites where it could be used to scan patients who require constant medical attention. Perhaps most interestingly, the high level of portability and the low infrastructure requirements of our IM system make it ripe for deployment on a mobile platform (i.e., an OPM-MEG system in a truck). This would have multiple uses – e.g., it could be deployed as a facility that could visit multiple epilepsy clinics – enabling patients at multiple sites to benefit from MEG (Rampp et al., 2019) whilst minimising cost. It could also enable deployment of OPM-MEG in “field trials”, for example at military training establishments or sports grounds to monitor concussion (Rier et al., 2021). Finally, a mobile platform could be used to gather large data sets from multiple geographic locations – including varying socioeconomic regions – something that is challenging when scanners are based in universities. In sum, the highly portable nature of the IM system offers new opportunities that are not accessible using conventional (cryogenic) MEG technology.

Figure 6 shows that our IM system can be used simultaneously with EEG to gather multi-modal electrophysiological data. This is not the first time OPM-MEG/EEG has been carried out (e.g., (Boto et al., 2019; Feys, et al., 2023; Seedat et al., 2023)), but it is noteworthy that it is the first simultaneous OPM-MEG/EEG recording with closed-loop operation and this mode (which requires continual operation of on-board-sensor electromagnetic coils) does not appear to impact significantly the quality of the EEG data. Our aim in this study was simply to demonstrate that concurrent recordings are possible. This is important for future clinical use since the acquisition of multi-modal data means clinical neurophysiologists can acquire EEG – which they are highly familiar with – at the same time as OPM-MEG data, which offers significant advantages in terms of spatial accuracy. As such, it will allow clinicians to relate EEG features which are detected from specific EEG montages to the high density, source localised OPM-MEG signals which will provide a bridge for translation of this new technology into the clinic. It was not our intention to directly compare OPM-MEG and EEG; nevertheless anecdotally, results from both modalities show clear beta and gamma band responses, even during head movement. However, whereas OPM-MEG consistently placed the gamma response in primary visual cortex, the EEG localisation placed it lower, in the cerebellum – this is likely a result of the difficulties with EEG caused by a poor model of the inhomogeneous conductivity profile of the head.

Finally, from a practical point of view, the IM system performed well. In previous OPM-MEG systems robustness has been a key concern – in particular the number of channels that are lost in a measurement. Here, across 32 experiments using our IM system we lost (on average) 3 ± 5 channels. Where we did lose channels, the reason was typically a connection between the sensor head and ribbon cable. Sensor heads are connected using a latch which clamps down on the ribbon cable, making an electrical connection. This necessitates minimal tolerance when manufacturing the cables, since even small changes in cable thickness can make the latch connector loose, and consequently the connection temperamental (this was also the likely reason for the marginally increased empty room noise in the IM system.) This is something that should be altered in future generations of this system. Despite this minor limitation, the IM system performed well. The set-up time for 64 sensors was typically around three minutes – this includes the time to heat the vapour cells and lasers, lock their temperature with PID controllers, and optimise all sensor parameters, zero the field within each cell, calibrate the sensor and turn on the closed-loop. Each OPM sensor head has slightly different properties meaning that control parameters must be optimised on a per sensor basis (much like superconducting quantum interference devices (SQUIDs) must each be individually tuned in a conventional MEG system). In the IM system, because these parameters are optimised and set on sensor start-up, sensor heads can be swapped easily with no requirement for anything other than a sensor restart following a swap. This is a significant practical advantage when running the system, adding further modularity to the design.

## CONCLUSION

We reported a fundamentally new OPM-MEG system design with miniaturised and integrated electronic control, a high level of portability, and significantly improved dynamic range. We have demonstrated that this instrumentation offers equivalent measures of induced and evoked neuro-electrical responses to stimuli compared to an established instrument, and that it offers improved dynamic range, up to and including 8 nT shifts in background field. We have shown that the system is effective in gathering data during participant movement (including a sitting-to-standing paradigm) and that it is compatible with simultaneous EEG recording. Finally, we demonstrated portability by moving the system between two laboratories. Overall, our new system represents a significant step forward for OPM-MEG and offers an extremely attractive platform for next generation functional medical imaging.

## ACKNOWLEDGEMENTS

This work was supported by an Engineering and Physical Sciences Research Council (EPSRC) Healthcare Impact Partnership Grant (EP/V047264/1). We acknowledge support from the UK Quantum Technology Hub in Sensing and Timing, funded by EPSRC (EP/T001046/1), and a Medical Research Council (MRC) Mid-Range Equipment grant (MC_PC_MR/X012263/1). Sensor development was made possible by funding from the National Institutes of Health (R44MH110288). O.F. is supported by the Fonds pour la formation à la recherche dans l’industrie et l’agriculture (FRIA, Fonds de la Recherche Scientifique [FRS-FNRS], Brussels, Belgium). P.C. is supported by the Fonds Erasme (Convention « Alzheimer », Brussels, Belgium). X.D.T. is a Clinical Researcher at the FRS-FNRS.

## CONFLICTS OF INTEREST

V.S. is the founding director of QuSpin, a commercial entity selling OPM magnetometers. J.O., D.B. and C.D. are employees of QuSpin. E.B. and M.J.B. are directors of Cerca Magnetics Limited, a spin-out company whose aim is to commercialise aspects of OPM-MEG technology. E.B., M.J.B., R.B., N.H. and R.H. hold founding equity in Cerca Magnetics Limited. HS, MR and ZT are employees of Cerca Magnetics Limited.

## DATA AND CODE AVAILABILITY

All data were acquired by the authors and are available upon request.

All code was custom developed in-house using MATLAB and is available from the authors on request.

## SUPPLEMENTARY INFORMATION

**Figure S1:**
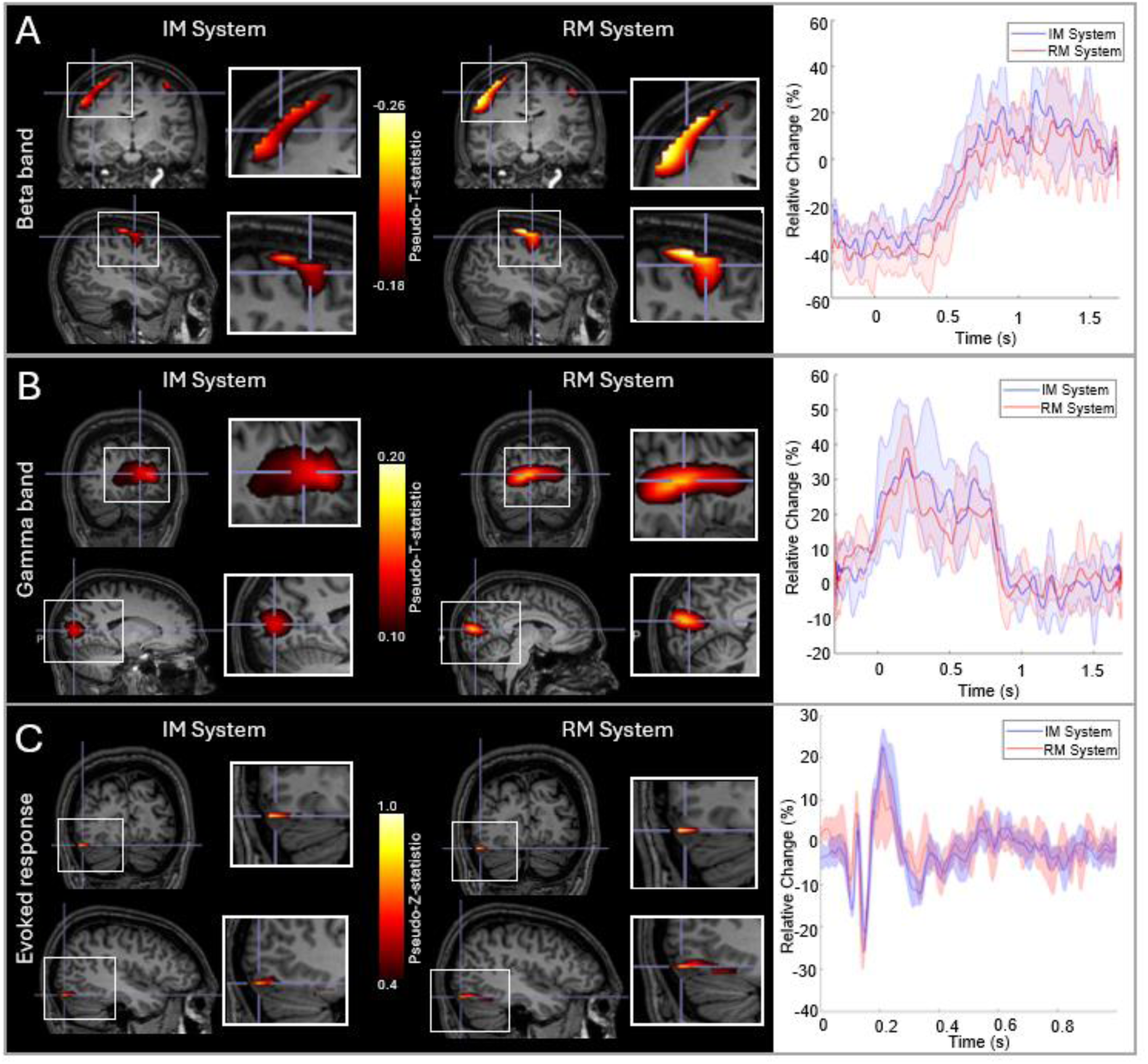
RM and IM system comparison for S1. Layout equivalent to Figure 2, but results shown for Subject 1 (S1).

**Figure S2:**
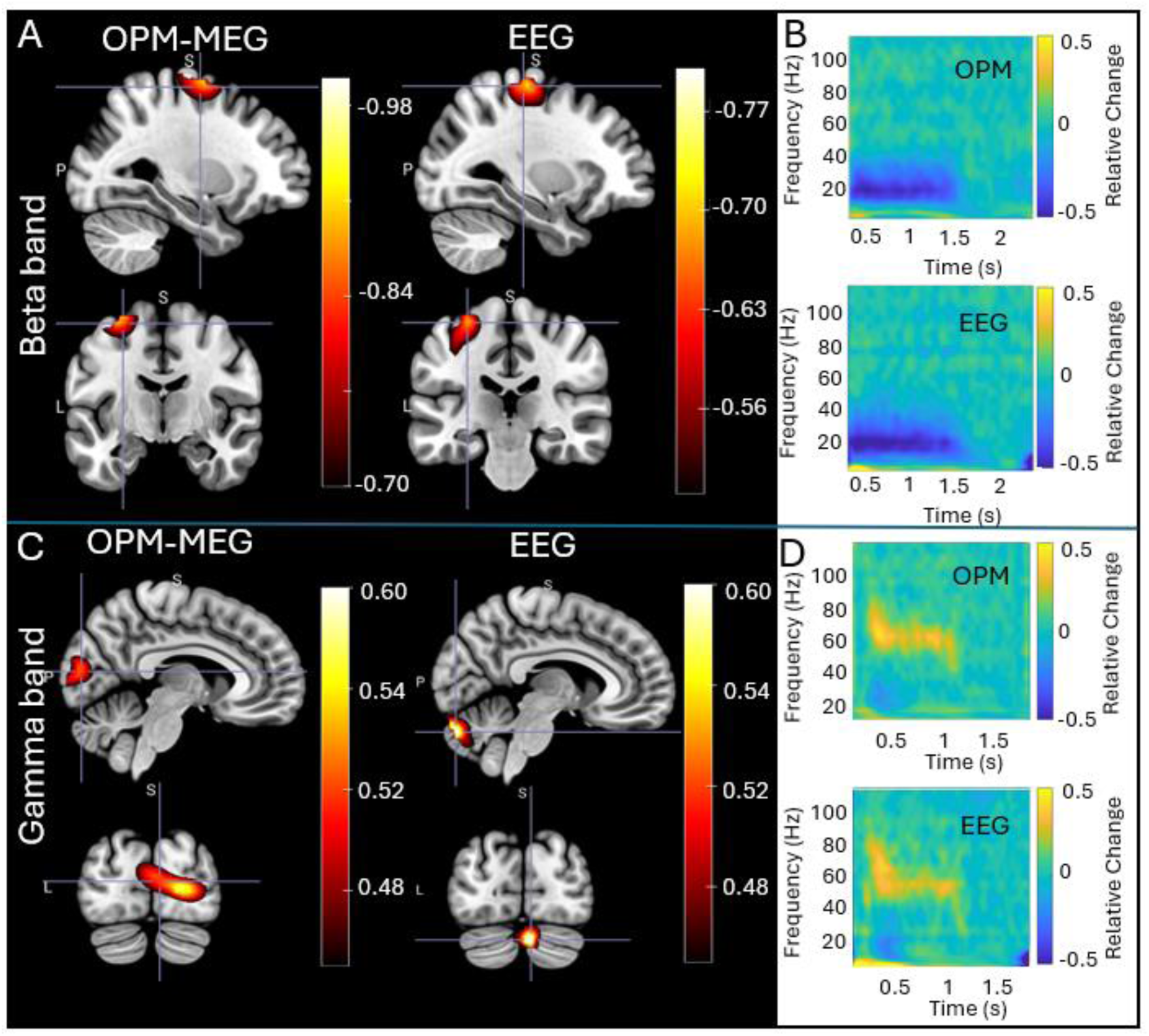
Concurrent OPM-MEG/EEG. Same as Figure 6 but in the static case (i.e. no head motion).

